# Autolysin-Independent DNA Release in *Streptococcus pneumoniae in vitro* Biofilms

**DOI:** 10.1101/322818

**Authors:** Mirian Domenech, Ernesto García

## Abstract

Biofilms are defined as layers of cells of microorganisms adhered to the surface of a substrate and embedded in an extracellular matrix and provide an appropriate environment for increased genetic exchange. Extracellular DNA (eDNA) is an essential component of the extracellular matrix of microbial biofilms, but the pathway(s) responsible for DNA release are largely unknown. Autolysis (either spontaneous or phage-induced) has been proposed the major event leading to the appearance of eDNA. The ‘suicidal tendency’ of *Streptococcus pneumoniae* is well-known, with lysis mainly caused by the triggering of LytA, the major autolytic amidase. However, the LytC lysozyme and CbpD (a possible murein hydrolase) have also been shown involved. The present work examines the relationship between eDNA, autolysins, and the formation and maintenance of *in vitro* pneumococcal biofilms, via fluorescent labelling combined with confocal laser scanning microscopy, plus genetic transformation experiments. Bacterial DNA release mechanisms other than those entailing lytic enzymes were shown to be involved by demonstrating that horizontal gene transfer in biofilms takes place even in the absence of detectable autolytic activity. It had been previously suggested that the quorum sensing systems ComABCDE and LuxS/AI-2 are involved in the production of eDNA as a response to the accumulation of quorum sensing signals, although our immunofluorescence results do not support this hypothesis. Evidence that the release of DNA is somehow linked to the production of extracellular vesicles by *S. pneumoniae* is provided.

## IMPORTANCE

Most human bacterial infections are caused by microorganisms growing as biofilms. Bacteria in biofilms are less susceptible to antimicrobials and to killing by the host immune system, are very difficult to eliminate and cause recalcitrant and persistent diseases. Extracellular DNA is one of the major components of the bacterial biofilm matrix. In the present study, we provide direct evidence of the existence of biologically active (transforming), extracellular DNA in *Streptococcus pneumoniae* biofilms. In previous studies, the involvement of three pneumococcal choline-binding proteins with autolytic activity (LytA, LytC and CbpD) in DNA release had been reported. In contrast, we demonstrate here that pneumococcal *in vitro* biofilms do contain eDNA, even in the absence of these enzymes. Moreover, our results suggest that DNA release in *S. pneumoniae* biofilms is connected with the production of extracellular vesicles and that this DNA is associated to the outer part of the vesicles.

The human pathogen *Streptococcus pneumoniae* is a leading cause of pneumonia, meningitis and bloodstream infections in the elderly, and one of the main pathogens responsible for middle ear infections in children. It is carried asymptomatically in the nasopharynx of many healthy adults, and in as many as 20-40% of healthy children (colonization begins shortly after birth) (1). Pneumococcal biofilms appear on adenoid and mucosal epithelium in children with recurrent middle-ear infections and otitis media with effusion, and on the sinus mucosa of patients with chronic rhinosinusitis, and they can be also formed *in vitro* (2, 3). Biofilm formation in *S. pneumoniae* is an efficient way of evading both the classical and the PspC-dependent alternative complement pathways (4).

Over 60% of all human bacterial infections, and up to 80% of those that become chronic, are thought to involve growth in biofilms. A biofilm is defined as an accumulation of microorganisms embedded in a self-produced extracellular matrix (ECM) adhered to an abiotic or living surface (5). The ECM is composed of different polymers, mainly polysaccharides, proteins, and nucleic acids. The requirement of extracellular deoxyribonucleic acid (eDNA) in ECM formation and maintenance has been documented in a variety of Gram-positive and Gram-negative bacteria (6). eDNA binds to polysaccharides and/or proteins, protecting bacterial cells from physical and/or chemical challenges, as well as providing biofilms with structural integrity. It is a major component of the *S. pneumoniae* ECM (7–11). Various pneumococcal surface proteins, *e.g.,* several members of the choline-binding family of proteins (CBPs) (11, 12) and the pneumococcal serine-rich repeat protein (PsrP) (13), form tight complexes with eDNA via electrostatic interactions, a mechanism proposed widespread among microorganisms (14). It should be noted that PsrP appears in ≈ 60% of clinical pneumococcal isolates, whereas the main CBPs — LytA (SPD_1737/SPD_RS09190), LytB (SPD_0853/SPD_RS04550), LytC (SPD_1403/SPD_RS07385) or CbpD (SPD_2028/SPD_RS10645) among others — are present in all *S. pneumoniae* strains.

The source of eDNA may vary across microorganisms and in part appears because of autolysis, phage-induced lysis, and/or active secretion systems, as well as through association with extracellular vesicles (EV). In *S. pneumoniae*, the release of DNA during limited lysis of the culture, *i.e.*, by controlled autolysis directed by the main CBP autolysins (LytA *N*-acetylmuramoyl-l-alanine amidase [EC 3.5.1.28; NAM-amidase] and LytC lysozyme [EC 3.2.1.17; muramidase]), as well as prophage-mediated lysis, have been proposed as biofilm-promoting in part of the bacterial population (9, 11). The NAM-amidase LytA, the main autolytic enzyme of *S. pneumoniae*, is kept under control by lipoteichoic acid — a membrane-bound teichoic acid that contains choline — during exponential growth (15), and regulated at the level of substrate recognition (16). LytC also acts as an autolysin when pneumococci are incubated at about 30°C, a temperature close to that of the upper respiratory tract (17); this lysozyme might be post-transcriptionally inhibited by CbpF (SPD_0345/SPD_RS01835) (18).

Studies in different microorganisms suggest that the appearance of eDNA in biofilms may also be a response to the accumulation of quorum sensing (QS - a cell density-dependent communication system that regulates cooperative behavior) signals (for a recent review, see reference 19). Two early studies on genetic transformation in planktonic cultures showed *S. pneumoniae* to release measurable amounts of DNA in the absence of detectable autolysis (20). Although these pioneering studies were forgotten for years, more recent investigations have shown that LytA, LytC and CbpD (a putative cell wall-degrading enzyme) are directly responsible for autolytic DNA release from only a subset of cells in competent pneumococcal planktonic cultures (21, 22). In fact, the killing of non-competent sister cells by competent pneumococci — a phenomenon named fratricide — promotes allolysis and DNA release (23). It is well known that competence induction in *S. pneumoniae* depends on a QS mechanism (24). LytA, LytC and CbpD are all required for allolysis (and for the concomitant DNA release) when pneumococci grow under planktonic conditions (25). It is currently believed that the limited damage caused by CbpD activates LytA and LytC, resulting in the more extensive lysis of target cells than that achieved by CbpD alone. LytA and LytC are constitutively synthesized by non-competent cells. However, while the expression of LytA increases during competence, LytC is not part of the competence regulon in *S. pneumoniae* (26). A slightly different situation has been proposed to occur in biofilms. The lysis of target (non-competent) cells in biofilms requires CbpD to act in conjunction with LytC, whereas LytA is not required for efficient fratricide-mediated gene exchange (26). In these experiments, however, a direct visualization of eDNA in the ECM was not reported. Interestingly, the transcription of both *lytA* and *cbpD* also appears to be regulated by the LusX/autoinducer-2 (AI-2) QS (27).

Via the use of an *in vitro* biofilm model system, the present work provides evidence that a small (but significant) proportion of biologically active, eDNA in pneumococcal biofilms is released into the medium by an alternative (or complementary) pathway to cell autolysis. It would appear that this occurs independent of the activity of LytA, LytC and CbpD and the QS systems (ComABCDE and LuxS/AI).

## RESULTS

### Visualization of eDNA in the pneumococcal biofilm

*S. pneumoniae* R6 biofilms grown for 5 h at 34°C in C+Y medium were stained with a combination of SYTO 59, DDAO (7-hydroxy-9H-[1,3-dichloro-9,9-dimethylacridin-2-one]) and anti-double-stranded (ds) DNA monoclonal antibodies (α-dsDNA). When examined under the confocal laser scanning microscope (CLSM), abundant, mostly cell-associated eDNA was observed (Fig. 1). However, when scanned at 488 nm (green), immunostained eDNA appeared as a lattice-like array consisting of long DNA fibers, mainly at the top of the biofilm (Fig. 1C). Only seldomly they were associated with the cocci (marked with yellow arrows in Fig. 1I). At the bottom of the biofilm, small areas of what appeared to be compacted eDNA (but no fibers) were seen (data not shown).

**FIG 1.**
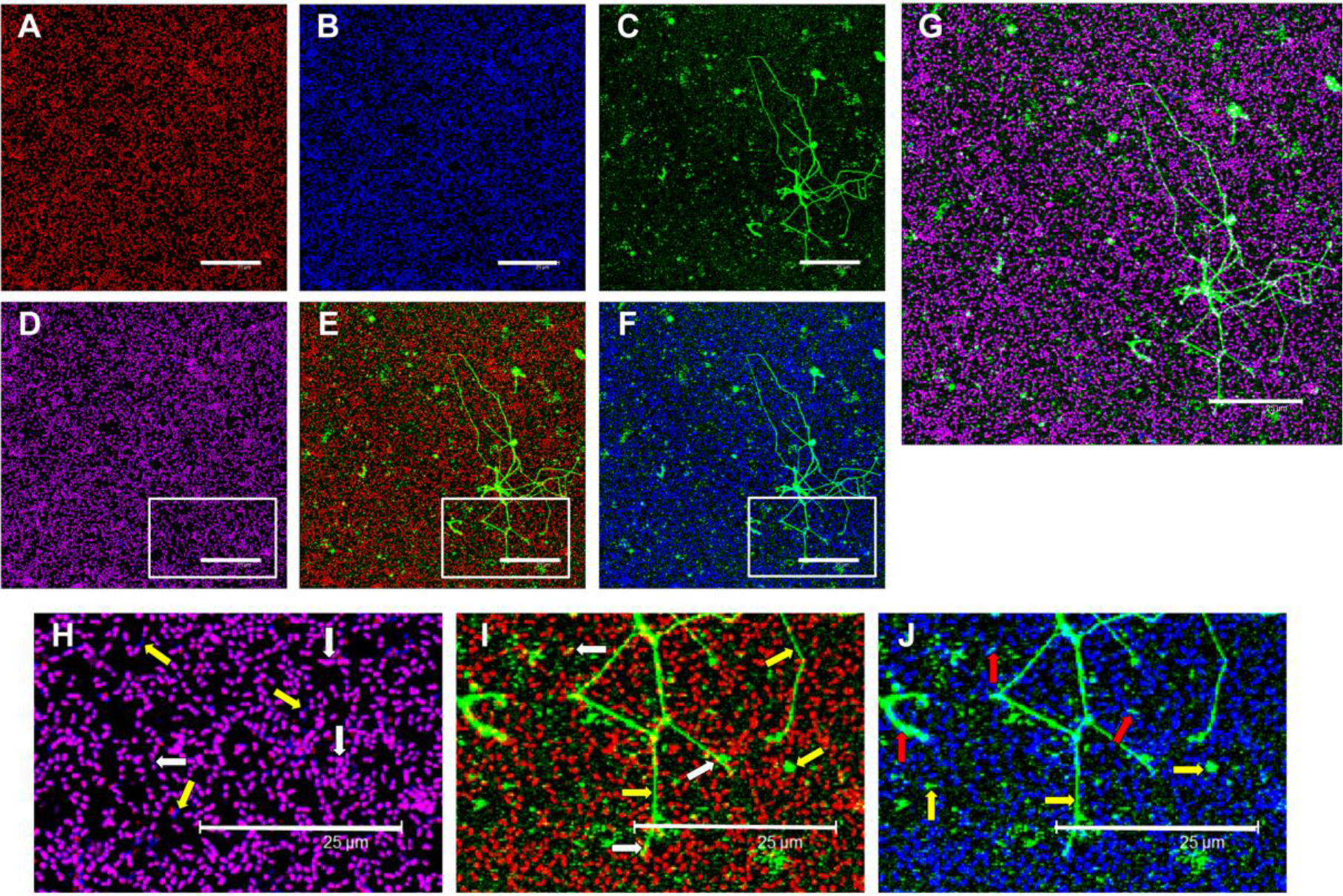
Evidence of the existence of eDNA in pneumococcal biofilms using CLSM. A biofilm of *S. pneumoniae* R6 grown for 5 h at 34°C in C+Y medium was stained with a combination of SYTO 59 (A, red), DDAO (B, blue), and α-dsDNA, followed by Alexa Fluor-488 goat anti-mouse IgG (C, green). Image D is a merge of channels A and B. Image E is a merge of channels A and C. Image F is a merge of channels B and C. Image G is a merge of the three channels. Images H, I and J correspond, respectively, to an enlarged vision of the area marked with squares in D, E and F. Yellow arrows point to eDNA stained only with DDAO or α-dsDNA-Alexa Fluor-488. White arrows indicate the location of SYTO 59-stained bacteria together with eDNA labelled either with DDAO (H, magenta) or with α-dsDNA-Alexa Fluor-488 (I, yellow). The red arrow in image J points to doubly labelled (DDAO plus α-dsDNA-Alexa Fluor-488) eDNA (J, light blue). Scale bars = 25 μm.

Notably, DNA fibers were not observed when DDAO was employed. Many reports have used DDAO for staining eDNA in biofilms (28). However, a recent evaluation of eDNA stains in biofilms of various species has shown that this compound was neither completely cell impermeant nor capable to reveal DNA-containing fibrilar structures (29). As our results were in agreement with these data, DDAO was not used in additional experiments.

To study the dynamics of eDNA release, strain R6 was incubated under biofilm-forming conditions for up to 5 h at 34°C. Immunostaining with α-dsDNA of sessile (adherent) cells revealed the existence of eDNA even at early incubation times (Fig. 2). At 3 h, eDNA filaments were visible, although the majority of eDNA appeared as dots and patches, which may represent different stages of condensation of DNA-protein aggregates, as previously suggested (12). In 5 h-old biofilms, eDNA threads were infrequent and mainly located at the top of the biofilm (see above). Previous studies have revealed the existence of a mature ECM (consisting of DNA, proteins and polysaccharides) at this time point (11). Immunostaining planktonic cultures (*i.e.*, non-adherent cells) revealed the existence of eDNA filaments that were more abundant in younger than in older cultures (Fig. 2).

**FIG 2.**
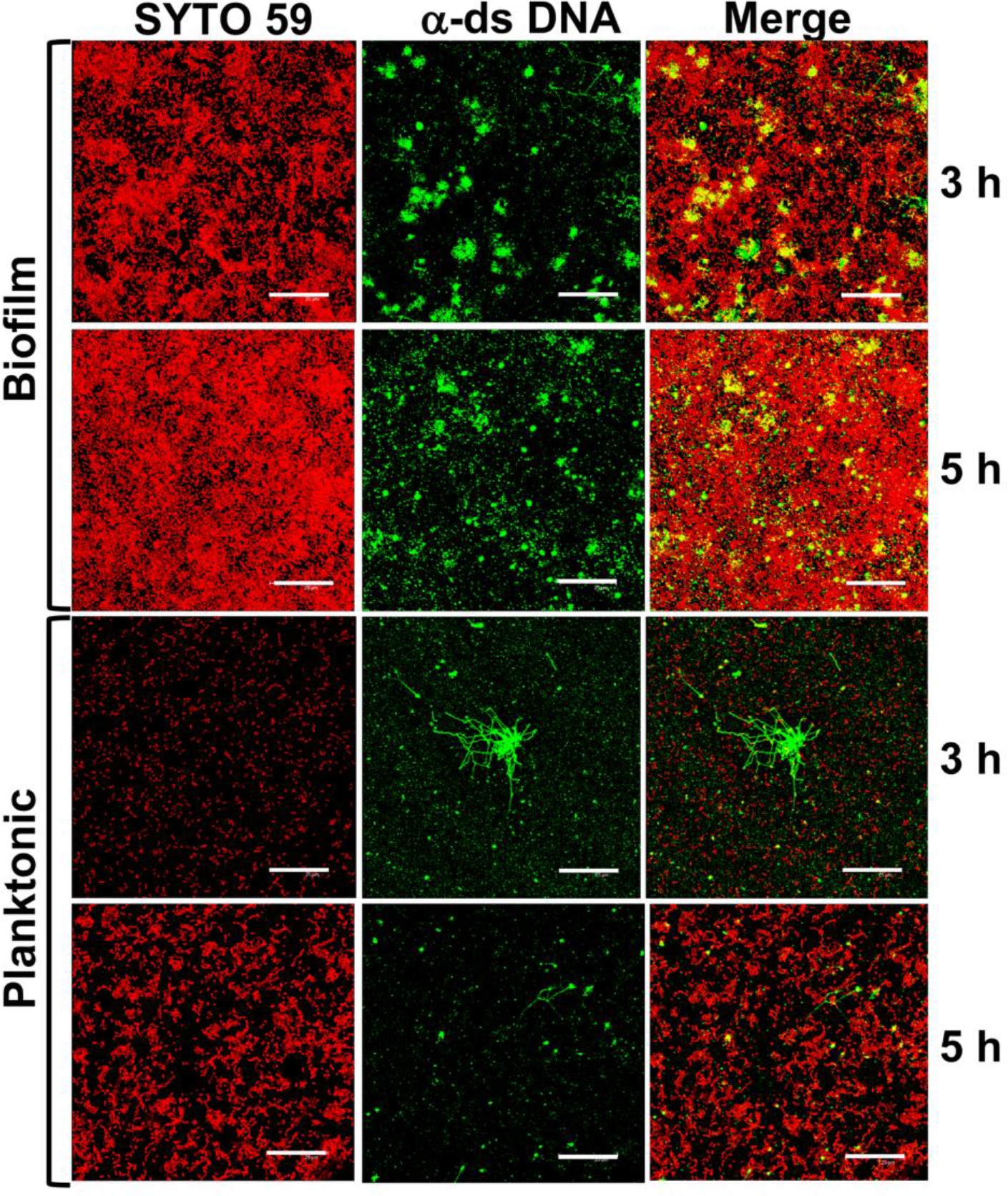
Dynamics of eDNA release in pneumococcal biofilms. *S. pneumoniae* R6 was grown under biofilm-forming conditions. After 3 and 5 h incubation at 34°C, adherent (biofilm) and non-adherent (planktonic) cells were independently stained with a combination of SYTO 59 (red) and α-dsDNA-Alexa Fluor-488 (green). Scale bars = 25 μm.

### Evidence of the presence of eDNA in biofilms formed by different pneumococcal mutants

The involvement of eDNA in biofilm formation and maintenance was ascertained using strain P046, which lacks the two main autolytic CBPs. Strain P234 was employed as a control; this has a point mutation in the *pspC* (= *cbpA*) gene (SPD_2017/SPD_RS10590), which encodes a CBP important in virulence (30). PspC, which lacks any autolytic activity and is partly involved in biofilm formation *in vitro* (on polystyrene microtiter plates) (7) and *in vivo* (in a murine nasopharynx colonization model) (3), also forms complexes with the eDNA (12). Compared to the parental R6 strain, both mutants showed reduced biofilm-forming capacity, in agreement with previous findings (7). Interestingly, biofilm formation, but not culture growth, was greatly impaired when pneumococci were grown in the presence of DNase I (Fig. 3). Moreover, the incubation of preformed biofilms with DNase I drastically diminished the number of biofilm-associated sessile cells. As a whole, however, preformed biofilms were less reduced by DNase I treatment than growing biofilms, strongly suggesting that eDNA is more important and/or more exposed during the early stages of biofilm formation. Alternatively, eDNA may become resistant to DNase enzymes during biofilm maturation by forming complexes with other macromolecules such as CBPs.

**FIG 3.**
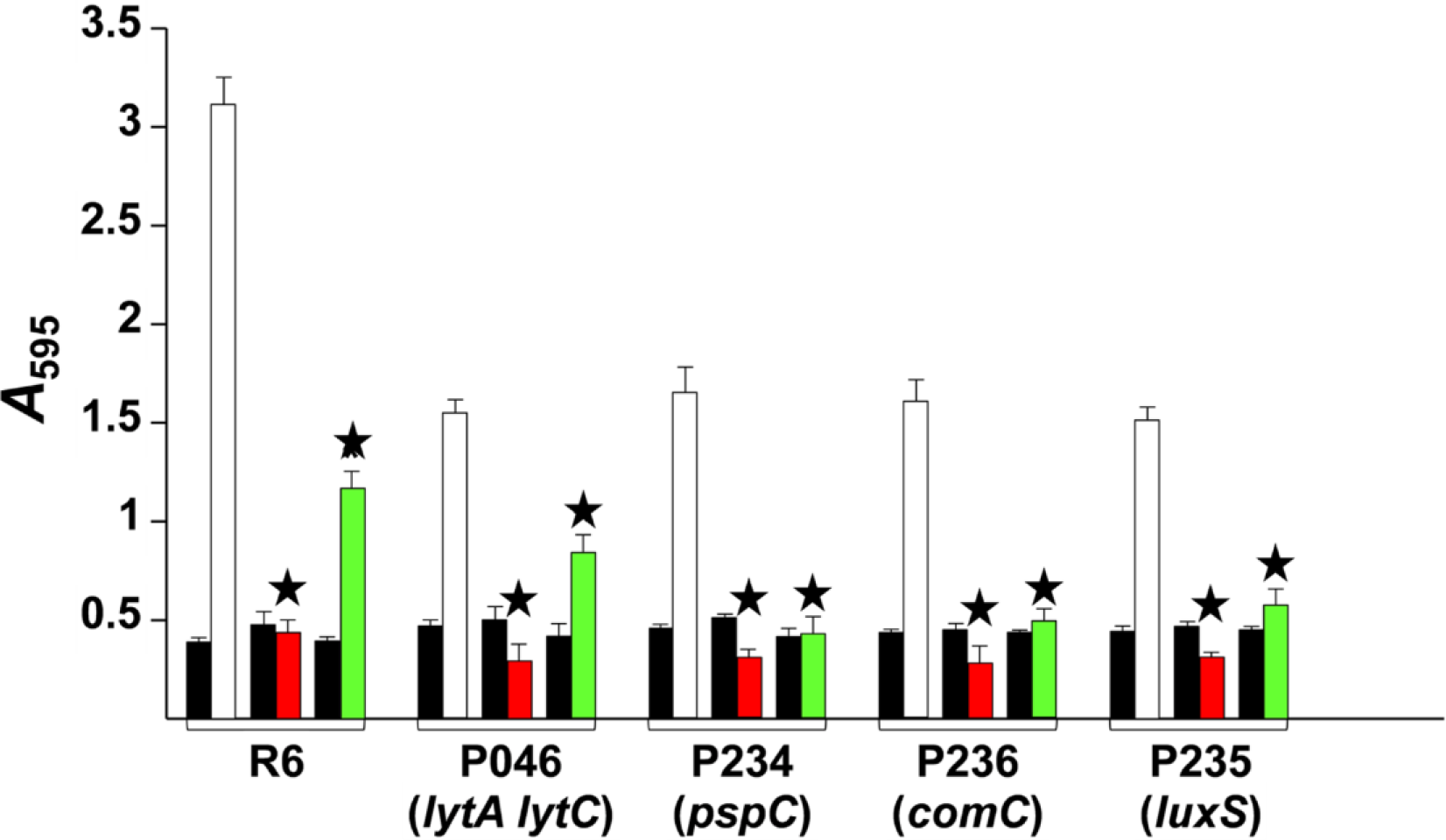
Inhibition and dispersal of pneumococcal biofilms with DNase I. (A) The indicated *S. pneumoniae* strains were grown overnight at 37°C to an *A*_550_ value of 0.5 (corresponding to the late exponential phase of growth) in C+Y medium, centrifuged, and adjusted to an *A*_550_ of 0.6 with fresh medium. The cell suspensions were then diluted 100-fold, and 200 μl aliquots distributed into the wells of microtiter plates, which were then incubated for 5 h at 34°C (open bars). Other samples received DNase I (100 μg ml^−1^) (red bars) and were incubated as above (inhibition assay). In other cases, and after biofilm development (4 h at 34°C), DNase I (green bars) was added at 100 μg ml^−1^, and incubation allowed to proceed for an additional 1 h at 34°C before staining with CV to quantify biofilm formation (dispersal assay). In all assays, black bars indicate growth (adherent plus non-adherent cells). *, *P* < 0.001 compared with the corresponding, untreated control.

The presence of eDNA in the biofilms of Δ*comC* (SPD_2066/SPD_RS10845) or Δ*luxS* (SPD_0309/SPD_RS1650) mutants was also analyzed. These genes are essential for the functioning of two QS systems documented as being involved in biofilm formation (see above). The *comC* gene codes for the pre-CSP (competence-stimulating peptide), which is matured and exported by the ComA-ComB complex as an unmodified 17-residue-long peptide pheromone. LuxS is an S-ribosylhomocysteine lyase, and is responsible for the production of the QS molecule homoserine lactone autoinducer 2 (AI-2). It has been reported that transcription of competence genes (including *lytA* and *cbpD*) is reduced in a Δ*luxS* strain (27). As shown above for two CBP mutants, the Δ*comC* and Δ*luxS* mutants produced only ≈50% of the biofilm formed by the R6 strain (Fig. 3). Positive evidence for the involvement of eDNA in formation and maintenance of these biofilms was also obtained by treatment with DNase I (Fig. 3).

### Contribution of lytic CBPs to eDNA release

The presence of eDNA in the ECM of biofilms formed by strain P046 — a double LytA^−^ LytC^−^ mutant — was unexpected since it is generally believed that at least one of the autolysins is required for DNA release (see above). To gain further insight, the existence of eDNA in the biofilm formed by a mutant deficient in CbpD (or its combination with the *lytA* and *lytC* mutations) was analyzed. A *pspC* mutant (strain P234) was included as a control. Notably, eDNA was present even in the ECM of strain P204, a mutant deficient in the three CBPs, *i.e.*, LytA, LytC and CbpD (Fig. 4). The morphology of the biofilms formed by strain P204 differed from those of R6, with the former biofilms containing fewer microcolonies and larger eDNA patches than the wild type.

**FIG 4.**
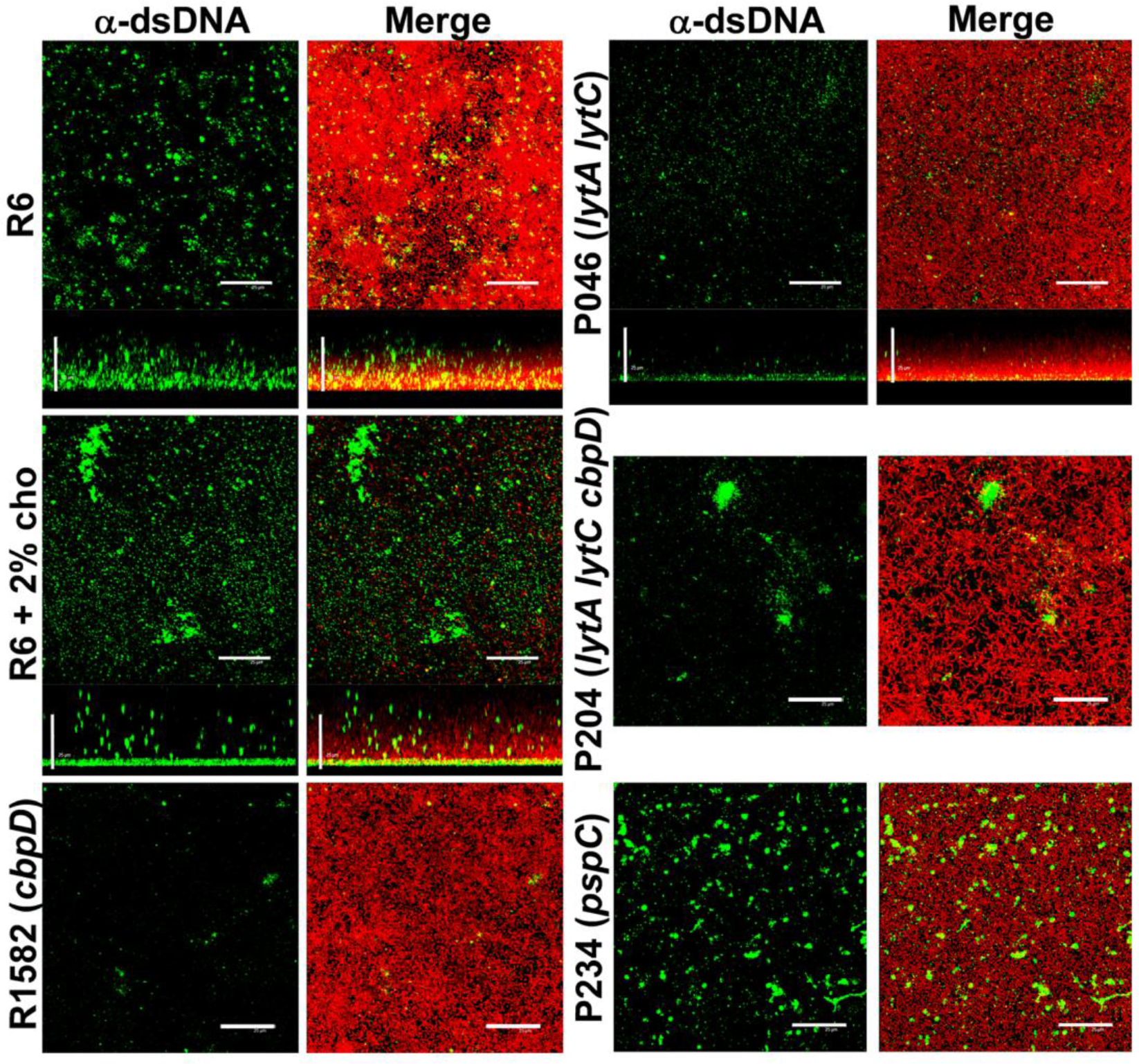
Influence of autolysins on biofilm formation and eDNA release revealed by CLSM. Biofilms were stained with a combination of SYTO 59 (red), and α-dsDNA, followed by Alexa Fluor-488 goat anti-mouse IgG (green). The R6 strain was also incubated in C+Y medium containing 2% choline chloride (R6 + 2% cho). Horizontal and vertical three-dimensional reconstructions of 55 (*x-y* plane) or 65 scans (*x-z* plane) are shown. Scale bars = 25 μm.

It is well known that when pneumococcal cells are incubated with 2% choline chloride, CBPs are released into the medium. Those with enzymatic activity are completely inhibited (31), but transformability is not altered (32). Under these conditions, however, the biofilm-forming capacity is severely reduced (7). This was confirmed in the present work, and might be attributed (at least in part) to the drastically diminished eDNA content of the biofilm. Quite unexpectedly, R6 biofilms formed in the presence of 2% choline chloride still showed the presence of eDNA (Fig. 4).

### The competence QS system is not involved in eDNA release

The presence of eDNA in biofilms formed by additional *S. pneumoniae* strains possessing combinations of mutations affecting the *comA* gene and various lytic genes was studied by CLSM. These strains were R391, P203, P204, and P213. The P147 strain (*ciaH*_Tupelo_VT_; SPD_0702/SPD_RS0372) was also included since in previous work our group has shown that CiaR/H, a two-component signal transduction system that mediates the stress response, is in some way: a) implicated in the triggering of the LytA autolysin (33), b) required for efficient *in vitro* biofilm formation and nasopharyngeal colonization in a mouse model (34), and c) involved in the control of competence for genetic transformation (35). CLSM observations of *in vitro* biofilms showed eDNA to be also present in the biofilms of every mutant tested (Fig. 5).

**FIG 5.**
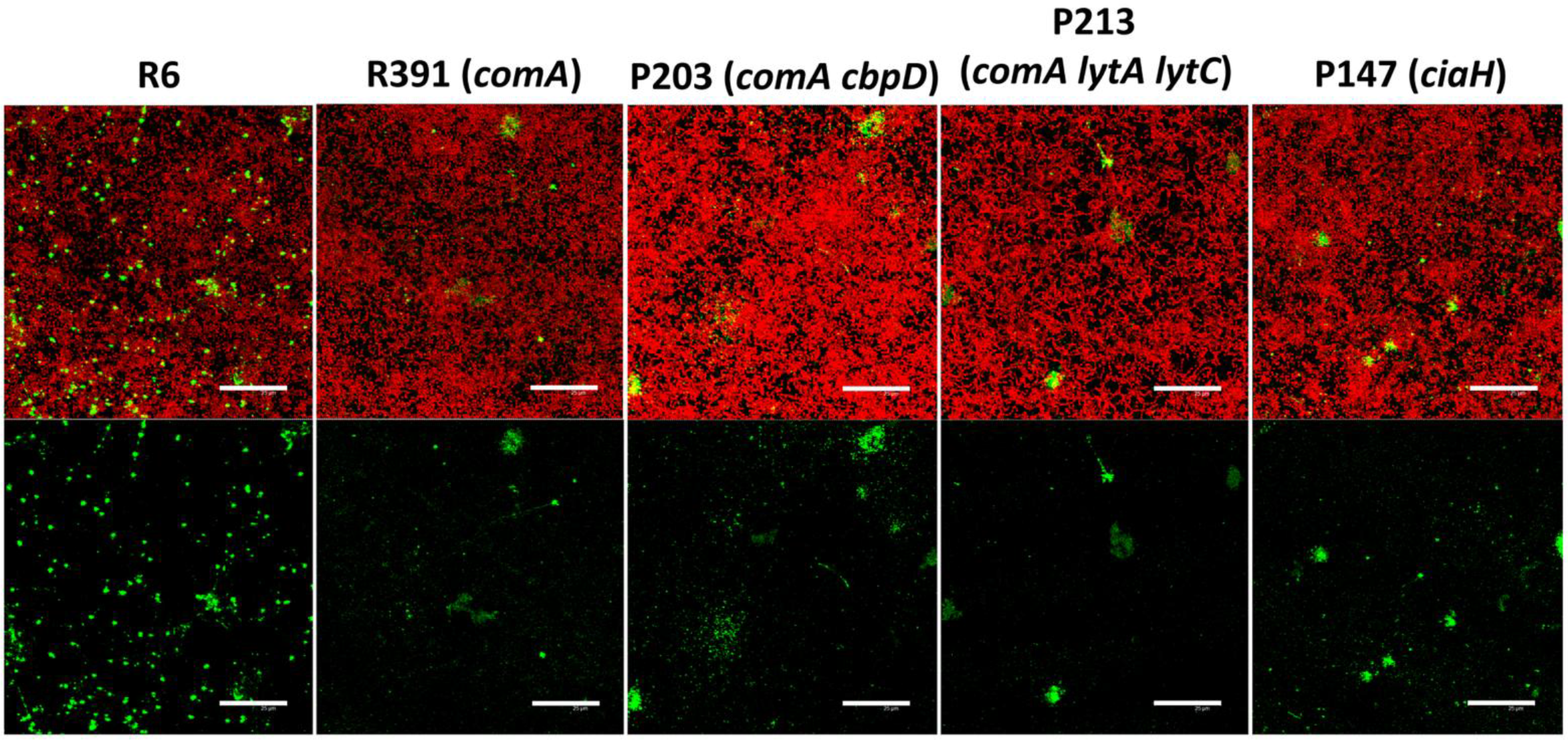
Influence of competence induction on biofilm formation and eDNA release revealed by CLSM. Biofilms were stained with a combination of SYTO 59 (red), and α-dsDNA, followed by Alexa Fluor-488 goat anti-mouse IgG (green). Scale bars = 25 μm.

### Transforming capacity of eDNA in pneumococcal biofilms

The above results indicate that *in vitro* pneumococcal biofilms are capable of releasing eDNA in a manner apparently independent of the activity of the three pneumococcal autolysins LytA, LytC and CbpD — even though these were generally believed necessary for DNA release in planktonic cultures. To investigate whether the eDNA in the biofilms formed by pneumococci simultaneously deficient in these three lytic enzymes has transforming activity, the efficacy of gene transfer in mixed biofilms was determined. For this, the reciprocal (donor and recipient) transforming capacity of two strain pairs was measured: in addition to being autolysin proficient (or not), one pair harbored the well-known low efficiency (LE) *nov1* marker (a C:G to T:A transition mutation conferring novobiocin resistance) (36), and the other the high efficiency (HE) marker *str41* (an A:T to C:G transversion), which bestows streptomycin resistance (37). To allow for further characterization of the direction of DNA transfer, the latter pair of strains used was also resistant to optochin. In pneumococcal transformation, the LE markers return 5-10% as many transformed cells as do HE markers (which are little degraded, or not at all). Table 1 shows that, in the biofilms, the spontaneous transformation of autolysin-proficient strains (P233 and P273) took place at levels typically observed in planktonic cultures when using naked chromosomal DNA (between 0.1 and 1% of total viable cells). Moreover, the heteroduplex DNA base mismatch repair system (Hex), which is responsible for marker-specific variations in transforming efficiency in planktonic cultures (38), was functional in the pneumococcal biofilms, as deduced from the relative transfer efficiency of the LE *nov1* and HE *str41* mutations (Table 1). In agreement with that reported by other authors (26), the transformation frequency was reduced by more than 100-fold in mixed biofilms formed by the mutants deficient in the three lytic enzymes. On average, however, one among 10^5^ pneumococci showed transformation to streptomycin resistance, demonstrating that a small amount of biologically active eDNA was still present in the ECM of the biofilms formed by the triple deficient mutant.

**TABLE 1.** Transformation efficiency in *S. pneumoniae* biofilms^a^

| Strain |  | Transformation frequency<sup>b</sup> |  |
| --- | --- | --- | --- |
|  |  | Str<sup>r</sup> | Nov<sup>r</sup> |
| P233 (Nov<sup>r</sup>) | P273 (Opt<sup>r</sup> Str<sup>r</sup>) | 0.45 ± 0.10 | 0.04 ± 0.02 |
| P272 (*lytA lytC*<br>*cbpD* Nov<sup>r</sup>) | P271 (*lytA lytC*<br>*cbpD* Opt<sup>r</sup> Str<sup>r</sup>) | $0.27 \times 10^{-2} \pm 0.05 \times 10^{-2}$ | $0.30 \times 10^{-3} \pm 0.19 \times 10^{-3}$ |

<sup>a</sup>Exponentially growing cultures of the indicated donor (and recipient) strains were mixed in polystyrene microtiter plates and incubated in C+Y medium at 34°C. After 4.5 h incubation, non-adherent cells were removed and biofilm-grown cells were washed and resuspended with PBS, diluted, and plated on blood agar plates containing Nov plus Str. Nov<sup>r</sup> Str<sup>r</sup> transformants were then picked from plates containing Opt to ascertain their parental strain. Total viable cells of each strain were determined using plates containing either Nov or Opt plus Str, and ranged from $1.5 \times 10^8$ to $1.8 \times 10^8$ for P233 + P273 biofilms, and from $3.2 \times 10^8$ to $3.4 \times 10^8$ CFU ml<sup>-1</sup> for P272 + P271 biofilms.
<sup>b</sup>Transformation frequency is defined as the number of transformants (CFU ml<sup>-1</sup>) multiplied by 100, divided by the total number of bacteria (CFU ml<sup>-1</sup>). Values correspond to the means ± standard errors for four independent experiments.

Previous studies performed with different bacteria have suggested that, in addition to autolytic phenomena, eDNA might be associated with EV. Recent evidence indicates that *S. pneumoniae*, actively sheds extracellular nano-sized EV, as do many other microorganisms (39–41). Whether pneumococcal EV contains DNA, however, was never examined. Here, high-speed sediments of biofilm filtrates — a crude preparation of EV with associated DNA (see Fig. S2 in the supplemental material)— were used to perform additional transformation experiments. The results confirmed a measurable amount of transforming DNA to be present in these crude preparations, and that this DNA could be destroyed by treatment with DNase I (Table 2). Comparable results were obtained when strain P271 was incubated under planktonic conditions.

**TABLE 2.**
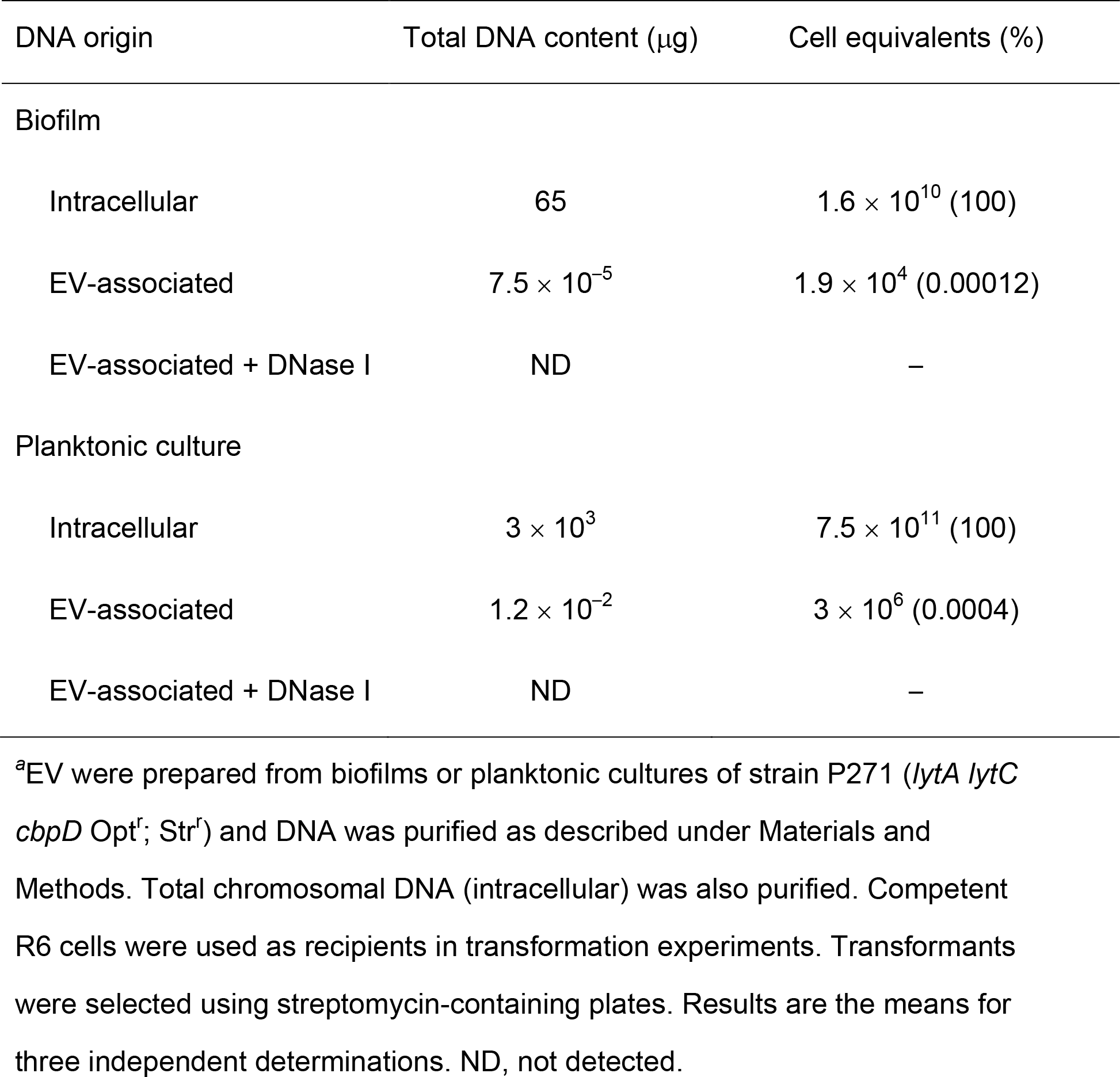
Transforming eDNA in pneumococcal biofilms and planktonic cultures^a^

## DISCUSSION

It has been known for decades that biofilm ECM contains eDNA, but its active role in biofilm appearance and maintenance was not recognized until Whitchurch *et al.* added DNase I to a *Pseudomonas aeruginosa* biofilm and watched the biofilm disappear (42). Since then, many reports have confirmed that a plethora of bacteria require eDNA to establish and maintain biofilms (6). The destruction of this eDNA provides a way of fighting biofilm-producing pathogens. Indeed, numerous clinical studies have shown that aerosolized DNase I (dornase alpha, a recombinant DNase I from the human pancreas) is highly effective in this respect, improving the lung function of patients with cystic fibrosis (43).

*In vivo* studies have shown that eDNA is a major element of biofilms. However, whether it is of bacterial or environmental (including host) origin (or both) is controversial; certainly it is difficult to determine which is the case under *in vivo* conditions. Studies using *in vitro* biofilm models (grown on plastic, glass, or other abiotic surfaces) provide a convenient way to screen large sets of strains, treatments, or growing conditions (44).

Among the possible origins of eDNA in pneumococcal biofilms (14), autolysins (either from bacterial or phage origin) have been proposed to have critical role. Previous studies performed at our own and other laboratories have employed indirect (*e.g.*, DNase I treatment and/or intrabiofilm transformation experiments) and direct methods (*e.g.*, staining with a variety of DNA-specific fluorophores) to disclose the presence of eDNA in pneumococcal biofilms (7–9, 11, 26, 27). Immunostaining with α-dsDNA combined with CLSM also revealed the existence of long filaments of eDNA in biofilms formed either by *Enterococcus faecalis* (45) or *Haemophilus influenzae* ECM (46). Moreover, the presence of eDNA-containing fibrous structures in the ECM of *Staphylococcus aureus* and *Propionibacterium acnes* biofilms has also been reported using α-dsDNA and atmospheric scanning electron microscopy (47, 48). The use of different technical approaches reinforces the idea that the fibrous assemblies of eDNA observed in some bacterial biofilms actually exist and also offers a novel perspective of the ECM structure of pneumococcal biofilms. Since most of the DNA filaments were found at the top of the mature biofilm (where actively growing cells reside) within 3 h of growth (Fig. 1), it is unlikely that these eDNA fibers were formed exclusively via autolysis. The same conclusion might be drawn from the CLSM images of immunostained planktonic cultures (Fig. 2).

In agreement with previous results, pneumococcal mutants either lacking the major autolysins or deficient in Com or LuxS/AI2 QSs showed limited biofilm-forming capacity. However, the results obtained following DNase I treatment of growing or pre-formed biofilms strongly suggest that those biofilms still contain eDNA. Direct CLSM visualization of the corresponding immunostained biofilms fully confirmed the presence of eDNA, even when the pneumococci were incubated in the presence of 2% choline chloride; this is known to induce the complete non-competitive inhibition of CBPs with enzyme activity, and to release all CBPs, whether enzymatic or not, from their bacterial surface attachments. It should be noted, however, that even at this high concentration, choline does not separate the CBP-DNA complexes that form part of the ECM of *S. pneumoniae* biofilms (11, 12). Nevertheless, since the complete inhibition of the enzymatic activity of autolysins takes place under these conditions, an exclusive, direct role for such enzymes in eDNA release appears to be unlikely. More direct evidence was obtained using biofilms formed by strain P204, a triple Δ*lytA* Δ*lytC* Δ*cbpD* mutant, in which the presence of eDNA was also verified (Fig. 4). The release of eDNA was also observed in biofilms formed by Δ*comA* mutants or combined Δ*comA*/autolysin-deficient mutations. Most notably, the existence of biologically active eDNA in pneumococcal biofilms — even when autolysin-deficient strains were used — was fully confirmed by *in situ* reciprocal transformation experiments (Table 1). Deletion of either *lytA*, *lytC* and/or *cbpD* does not alter the transformability of the mutants compared to their parental strains when chromosomal DNA or plasmid(s) is used as donor material (17, 49). Wei and Håvarstein studied the impact of LytA, LytC and CbpD on transformation efficiency in biofilm-grown pneumococci using an approach with some similarity to that employed here (26). The authors made use of *in vitro* mixed biofilms containing spectinomycin-resistant (Spc^r^) Δ*comA* ‘attacker cells’ — harboring concomitant deletions in the three autolysin genes (strain SPH149), or not (strain RH1) — and Nov^r^ non-competent (Δ*comA* Δ*comE*) ‘target cells’ (strain RH401). Upon the addition of CSP, the attackers (but not the targets, due to their being Δ*comE*) acquired competence and were transformed by the DNA released from the target cells through the fratricidal killing caused by the lytic enzymes induced by the attackers. A *ca.* 40-fold reduction in transformation efficiency was observed in biofilms composed of SPH149 attackers (autolysin-deficient) and RH401 cells (0.009%) with respect to that seen in biofilms containing the same target cells but involving a lysis-proficient strain (RH1) as the attacker (0.34%). Moreover, a further near 40-fold reduction in gene transfer frequency (0.00026%) was seen when SPH149 attackers were incubated with Δ*lytA* Δ*lytC* targets cells (strain SPH148) (26). Target cells containing a Δ*cbpD* mutation were not tested, possibly because, as previously reported for mixed planktonic cultures, CbpD-proficient attacker cells (RH1) were 1000-fold more efficient in transformations involving *cbpD*^+^ target cells than were CbpD-deficient attacker cells (49).

Pneumococcal biofilms that form during nasopharyngeal colonization may provide an optimal environment for increased genetic exchange with enhanced natural transformation *in vivo* (3). The presence of eDNA in the biofilm matrix has generally been attributed to the autolysis of a subpopulation of cells via fratricidal killing, suicidal killing, and/or the controlled release of DNA via signal transduction (19). Autolysis-independent eDNA release has been documented in some Gram-positive bacteria including *Bacillus subtilis* (50), enterococci (45, 51), and staphylococci (52, 53). Taking the present results together, it is clear that mechanisms involved in active eDNA release (perhaps associated with the production of EV), other than those directly dependent on autolysins, are at work in *S. pneumoniae.* This is important since an important feature of biofilms is the development of chemical gradients (i.e., pH, redox potential, and ions) (reviewed in reference 28). For example, in *P. aeruginosa* biofilms the pH value towards the center of a microcolony (≈ 6.0) is lower than that at the edge of the biofilm or in the growth medium (≈ 6.8) (54). Although not directly tested, a similar situation might be relevant in pneumococcal biofilms because it is well known that *S. pneumoniae* autolysis is inhibited at low pH values (≤ 6.0) (55). In this case, autolysin-independent DNA release would allow the maintenance of horizontal gene transfer events within the biofilm. Although the amount of biologically active DNA released in the absence of detectable autolytic activity is limited, it appears to be sufficient to partially promote a biofilm lifestyle and, importantly, to allow horizontal gene transfer. It is also possible that, as suggested by the CLSM observations, a substantial amount of eDNA may be present in biofilms, although lacking most transforming activity. It is tempting to speculate that the DNA-protein complexes present in the ECM may hinder the biological activity of eDNA as a transforming agent while contributing to the formation and preservation of the biofilm.

As reported for *Streptococcus mutans*, lysis-independent EV can transport DNA, contributing to horizontal gene transfer (56). Recent results also suggest that signal transduction mechanisms may be involved in the regulation of EV production in some Gram-positive bacteria (57, 58), but evidence for this is lacking in *S. pneumoniae*. As strongly suggested by the inhibitory effect of DNase I treatment in transforming efficiency, the biologically active eDNA released independent of detectable autolysis in *S. pneumoniae* appears to be located outside the EV. Although the loading of DNA into EV is thought to be widespread, experimental evidence showing that most EV-associated genomic DNA is present externally has been recently shown in *P. aeruginosa* (59, 60). However, other possibilities are also conceivable. For example, the stability of EV under various conditions may vary and a vesicle that loses membrane integrity may ‘leak’ its constituents into the supernatant (vesicle destabilization), rather than break up, as reported in *Bacillus anthracis*, *S. aureus* and other microorganisms (61). Interestingly, a very recent study has reported that the LytA NAM-amidase is not needed for production of EV, although the presence of EV-associated DNA was not analyzed (41).

An alternative (or complementary) mechanism for eDNA release may be bacterial type IV secretion systems (T4SS) that selectively deliver macromolecules to other cells or to the extracellular medium. An outstanding feature of these secretion systems is their ability to secrete both proteins and DNA molecules, a particularity that distinguishes them from other types of secretion system. The existence of a type IV secretion-like system involved in eDNA secretion was first described in *Neisseria gonorrhoeae* (62). More recently, it has been found that the release of eDNA from the cytoplasm of *H. influenzae* into the ECM requires the expression of an inner-membrane complex with homology to type IV secretion-like systems, plus the ComE outer-membrane pore through which the type IV pilus is extruded (63). Type IV pili are surface-exposed fibers that mediate many functions in bacteria, including locomotion, adherence to eukaryotic cells, biofilm formation, DNA uptake (competence), and protein secretion. Although initially considered to be exclusive to Gram-negative bacteria, they are also present in Gram-positive, although their role(s) is just beginning to emerge (64). Recently, several studies have revealed the existence of type IV competence-induced pili — predominantly composed of the ComGC pilin (SPD_1861; SPD_RS09815) — on the surface of *S. pneumoniae* cells (65). Since these pili bind DNA, it has been proposed that the transformation pilus is the primary DNA receptor on the bacterial cell during transformation in *S. pneumoniae*. It is tempting to speculate that, during competence, intracellular DNA may also be secreted into the ECM with the participation of the type IV pili, perhaps involving the aqueous pore formed by ComEC (SPD_0844; SPD_RS0450) in the cytoplasmic membrane (66). Further studies are required to test this hypothesis.

## MATERIALS AND METHODS

### Strains, media and growth conditions

Table 3 lists the pneumococcal strains used in this study; all were grown in pH 8-adjusted C medium (CpH8) supplemented with 0.08% yeast extract (C+Y) medium, or not, as required (7). Cells were incubated at 37°C without shaking, and growth monitored by measuring absorbance at 550 nm (*A*_550_). When used, antibiotics were added at the following concentrations: erythromycin 0.5 μg ml^−1^, kanamycin 250 μg ml^−1^, novobiocin 10 μg ml^−1^, optochin 5 μg ml^−1^, tetracycline 1 μg ml^−1^, spectinomycin 100 μg ml^−1^, and streptomycin 100 μg ml^−1^. DNase I (from bovine pancreas, DN25) was purchased from Sigma-Aldrich. For the construction of mutants (Table 3), the appropriate *S. pneumoniae* strains were transformed with chromosomal or plasmid DNA in C medium supplemented with 0.08% bovine serum albumin after treating cells with 250 ng ml^−1^ synthetic CSP-1 at 37°C for 10 min to induce competence, followed by incubation at 30°C during DNA uptake.

**TABLE 3.** *S. pneumoniae* strains used

| Strain | Description<sup>a</sup> | Source or reference |
| --- | --- | --- |
| R6 | Nonencapsulated D39 derivative; *lytA*<sup>+</sup> | 69 |
| R6BC | R6 *lytB::ermC lytC::tet*; Ery<sup>r</sup> Tet<sup>r</sup> | 7 |
| R800 | R6 derivative | 70 |
| M22 | A laboratory multi-resistant strain; *nov1 str41*; Nov<sup>r</sup>; Str<sup>r</sup> | 71 |
| M222 | M22 derivative; M22 transformed with *Streptococcus oralis* NCTC 11427 chromosomal DNA; Opt<sup>r</sup>; Str<sup>r</sup> | 72 |
| SPJV05 | D39 derivative *luxS* null mutant, Ery<sup>r</sup> | 73 |
| SPJV10 | D39 derivative *comC* null mutant, Ery<sup>r</sup> | 74 |
| P042 | R800 but *lytA::aphIII lytC::tet*; Kan<sup>r</sup> Tet<sup>r</sup> | 7 |
| P046 | R800 but *lytA::aphIII lytC::ermC*; Kan<sup>r</sup> Ery<sup>r</sup> | 7 |
| P104 | R6 but *lytA::cat*; Cm<sup>r</sup> | This study |
| P147 | R6 *ciaH*<sub>Tupelo_VT</sub> | 33 |
| P203 | R391 but *cbpD::spc*; R391 transformed with R1582 chromosomal DNA; Spc<sup>r</sup> | This study |
| P204 | P042 but *cbpD::spc*; P042 transformed with R1582 chromosomal DNA; Spc<sup>r</sup> | This study |
| P211 | P104 but *comA::kan*; P104 transformed with R391 chromosomal DNA; Kan<sup>r</sup> | This study |
| P213 | P211 but *lytC*; P211 transformed with R6BC chromosomal DNA; Tet<sup>r</sup> | This study |
| P233 | R6 transformed with M22 chromosomal DNA; Nov<sup>r</sup> | This study |
| P234 | R6 *pspC*; spontaneous mutant<sup>b</sup> | This study |
| P235 | R6 but *luxS* null mutant; R6 transformed with SPJV05 chromosomal DNA; Ery<sup>r</sup> | This study |
| P236 | R6 but *comC* null mutant; R6 transformed with SPJV10 chromosomal DNA; Ery<sup>r</sup> | This study |
| P271 | P204 transformed with M222 chromosomal DNA; Opt<sup>r</sup>; Str<sup>r</sup> | This study |
| P272 | P204 transformed with M222 chromosomal DNA; Nov<sup>r</sup> | This study |
| P273 | R6 transformed with M222 chromosomal DNA; Opt<sup>r</sup>; Str<sup>r</sup> | This study |
| R391 | R800 but *comA::kan*; Kan<sup>r</sup> | 25 |
| R1582 | R800 but *cbpD::spc*; Spc<sup>r</sup> | 25 |

<sup>a</sup>Cm, chloramphenicol; Ery, erythromycin; Kan, kanamycin; Nov, novobiocin; Opt, optochin; Tet, tetracycline; Spc, spectinomycin; Str, streptomycin; VT, vancomycin tolerance. <sup>r</sup>, resistant.
<sup>b</sup>The *pspC* mutation corresponds to a single base substitution (G:C to A:T) converting Trp (TGG) at position 512 to a stop codon (TAG). The wild type *pspC* gene encodes a 701 amino acid residue-long protein.

Biofilm formation was determined by the ability of cells to adhere to the walls and base of 96-well (flat-bottomed) polystyrene microtiter dishes (Costar 3595; Corning Incorporated), using a modification of a previously reported protocol (67). Unless stated otherwise, cells grown in C+Y medium to an *A*_550_ of ≈ 0.5-0. 6, sedimented by centrifugation, resuspended in an equal volume of the indicated pre-warmed medium, diluted 1/10 or 1/100, and then dispensed at a concentration of 200 μl per well. Plates were incubated at 34°C for 3, 4.5 and 5 h and bacterial growth determined by measuring the *A*_595_ using a VersaMax microplate absorbance reader (Molecular Devices). The biofilm formed was stained with 1% crystal violet (67).

### Intrabiofilm gene transfer

Exponentially growing cultures of donor strains (see above) were seeded together in a 1:1 ratio in polystyrene microtiter plates and incubated in C+Y medium at 34°C. Previous results have indicated that biofilm-grown pneumococcal cells must be actively growing to become competent (26). Hence, biofilms formation was allowed for only 4.5 h; nonadherent cells were removed and adherent cells were washed with PBS, disaggregated by gentle pipetting and slow vortexing (4). The latter were then serially diluted and plated on blood agar plates containing Nov plus Str. Nov^r^ Str^r^ transformants were then picked from plates containing Opt to ascertain their parental strain. For each donor strain the total number of viable cells was determined using blood agar plates containing either Nov or Opt plus Str.

### Preparation of extracellular vesicles and microscopical examination

EV-enriched centrifugation fractions were prepared from *S. pneumoniae* following standard procedures (61). Briefly, for biofilms, one liter of C+Y medium was inoculated with 10 ml of an exponentially growing culture of strain P271, distributed in 50 Petri dishes (10 cm diameter), and incubated at 34°C for 4.5 h. The non-adherent cells in the dishes were pipetted off and the biofilm-growing cells (6-7 × 10^9^ CFU) suspended in 75 ml of fresh C+Y medium. After centrifugation (9500 × *g*, 20 min, 4°C) the supernatant was filtered through a 0.2 μm pore-size filter (Millipore). The filtrate was centrifuged at 100,000 × *g* for 1 h at 4°C to sediment the vesicular fraction into a pellet. The supernatant was then discarded and the pellet suspended in a small volume of C+Y medium (EV fraction) and stored in aliquots at −20°C. Aliquots of the EV fraction were also treated with DNase I (10 μg ml^−1^) for 1 h at 37°C. EDTA (50 mM), SDS (1%) and proteinase K (100 μg ml^−1^) were then added and the mixtures incubated for 2 h at 37°C. Extraction was performed with phenol, and precipitation with ethanol, following standard procedures. Finally, the pellet was dissolved in a small volume of 10 mM Tris-HCl, pH 8.0 containing 1 mM EDTA. Cultures of the same pneumococcal strain were also grown under planktonic conditions until late exponential phase (≈ 3 × 10^8^ CFU ml^−1^) at 37°C without agitation, and processed as biofilm-grown pneumococci for EV preparation.

EV preparations (7 μl) were spotted on a glass slide, air dried, stained with the red fluorescence, membrane dye FM 4-64 (Molecular Probes), and incubated with α-dsDNA (ab27156, Abcam) followed by incubation with Alexa fluor 488-labeled goat anti-mouse IgG (A-11001, Invitrogen) (see below). Specimens were observed under a Leica DM4000B fluorescence microscope and viewed under a Leica HC PL APO63⨯/1.40-0.60 oil objective. *S. pneumoniae* P271 biofilms formed on glass surfaces were prepared for low-temperature scanning electron microscopy (LTSEM) as previously described (7), and the samples observed at −135°C using a DSM 960 Zeiss scanning electron microscope. For transmision electron microscopy (TEM) analysis, 5 μl of EV were placed for 5 min at room temperature on carbon-reinforced, Formvar-coated copper grids (300 mesh), which had been rendered hydrophilic by glow-discharge using a Quorum GloQube apparatus (Quorum Technologies). The sample was quickly washed with ultrapure water and negative staining was performed using 1% sodium phosphotungstate for 5 min. The excess stain was removed, and the sample was allowed to dry. Micrographs were taken on a JEOL JEM 1230 working at 80 kV.

### Quantification of eDNA

DNA quantification was performed by spectrophotometry (with concentrated samples) and/or using genetic transformation experiments (68). It is well known that a first-order concentration dependence (near unity) exists in chromosomal DNA transformation in pneumococci and other bacteria. This was confirmed in this study (see Fig. S1 in the supplemental material). It should be underlined that, since the size of the *S. pneumoniae* R6 genome is about 2 Mb, the DNA content of a single CFU, *i.e.*, a diplococcus, equals approximately 4 fg.

### Microscopic observation of biofilms

For the observation of *S. pneumoniae* biofilms by CLSM, pneumococcal strains were grown on glass-bottomed dishes (WillCo-dish) for 3-5 h at 34°C as previously described (11). Following incubation, the culture medium was removed and the biofilm rinsed with phosphate-buffered saline (PBS) to remove non-adherent bacteria. The biofilms were then stained with DDAO (2 μM) (H6482, Invitrogen), α-dsDNA (at 2-25 μg ml^−1^ each) and/or SYTO 59 (10 μM) (S11341, Invitrogen). All staining procedures involved incubation for 10-20 min at room temperature in the dark, except when biofilms were incubated with mouse α-dsDNA (2 μg ml ^−1^); this involved a fixation step at room temperature with 3% paraformaldehyde for 10 min. The biofilms were then rinsed with 0.5 ml PBS and incubated for 1 h at 4°C followed by 30 min incubation at room temperature in the dark with Alexa fluor 488-labelled goat anti-mouse IgG (1:500). After staining, the biofilms were gently rinsed with 0.5 ml PBS. Observations were made at a magnification of 63× using a Leica TCS-SP2-AOBS-UV or TCS-SP5-AOBS-UV CLSM equipped with an argon ion laser. Laser intensity and gain were kept the same for all images. Images were analyzed using LCS software from LEICA. Projections were obtained in the x-y (individual scans at 0.5-1 μm intervals) and x-z (images at 5-6 μm intervals) planes.

### Statistical analysis

Data comparisons were performed using the two-tailed Student *t*-test.

## ACKNOWLEDGMENTS

We thank J.-P. Claverys and J. Vidal for kindly providing pneumococcal strains, M. Moscoso, P. García and J. Yuste for helpful comments and critical reading of the manuscript, A. Burton for revising the English version, M. T. Seisdedos and G. Elvira for their help with CLSM, and E. Cano and S. Ruiz for skillful technical assistance.

This research was supported by grant SAF2017-88664 from Ministerio de Economía, Industria y Competitividad (MEICOM). Centro de Investigación Biomédica en Red de Enfermedades Respiratorias (CIBERES) is an initiative of the Instituto de Salud Carlos III (ISCIII).

## Supplemental Material

Additional information may be found in the online version of this article:

**FIG S1** Calibration curve for *S. pneumoniae*-transforming DNA. Competent R6 cells were used as recipient bacteria. Values represent the means ± standard errors for three independent transformation experiments. The dotted line corresponds to a slope of 1.

**FIG S2** Observation of EV produced by *S. pneumoniae* P271. (A-C) Fluorescent labeling of an EV preparation with FM 4-64 (red; A) and α-dsDNA, followed by incubation with Alexa fluor 488-labeled goat anti-mouse IgG (green; B). Image C is a merger of the two channels. Scale bars = 10 μm. (D-E) Electron micrographs of EVs. (D) LTSEM image of a pneumococcal biofilm. Arrows point to spherical blebs protruding from cells. The arrowhead indicates a putatively cell extruded EV. (E, F) TEM micrographs showing negatively stained EV. In some cases, EV appear to coalesce and fuse (F).

